# *In silico* mechanics of stem cells intramyocardially transplanted with a biomaterial injectate for treatment of myocardial infarction

**DOI:** 10.1101/2023.05.10.540185

**Authors:** YD Motchon, KL Sack, MS Sirry, NJ Nchejane, T Abdalrahman, J Nagawa, M Kruger, E Pauwels, D Van Loo, A De Muynck, L Van Hoorebeke, NH Davies, T Franz

**Affiliations:** Biomedical Engineering Research Centre, Division of Biomedical Engineering, Department of Human Biology, University of Cape Town, Observatory, South Africa; Cardiac Rhythm Management, Medtronic Inc, Minnesota, USA; Department of Biomedical Engineering, School of Engineering and Computing, American International University, Al Jahra, Kuwait; Cardiovascular Research Unit, University of Cape Town, Observatory, South Africa; Centre for X-ray Tomography, Department of Physics and Astronomy, Ghent University, Ghent, Belgium; XRE nv, Bollebergen 2B box 1, 9052 Ghent, Belgium; Bioengineering Science Research Group, Department of Mechanical Engineering, Faculty of Engineering and Physical Sciences, University of Southampton, Southampton, United Kingdom

**Keywords:** Biomaterial injection therapy, Cell mechanics, Cell therapy, Finite element method

## Abstract

**Purpose:** Biomaterial and stem cell delivery are promising approaches to treating myocardial infarction. However, the mechanical and biochemical mechanisms underlying the therapeutic benefits require further clarification. This study aimed to assess the deformation of stem cells injected with the biomaterial into the infarcted heart.

**Methods:** A microstructural finite element model of a mid-wall infarcted myocardial region was developed from *ex vivo* microcomputed tomography data of a rat heart with left ventricular infarct and intramyocardial biomaterial injectate. Nine cells were numerically seeded in the injectate of the microstructural model. The microstructural and a previously developed biventricular finite element model of the same rat heart were used to quantify the deformation of the cells during a cardiac cycle for a biomaterial elastic modulus (E_inj_) ranging between 4.1 and 405,900 kPa.

**Results:** The transplanted cells’ deformation was largest for E_inj_ = 7.4 kPa, matching that of the cells, and decreased for an increase and decrease in E_inj_. The cell deformation was more sensitive to E_inj_ changes for softer (E_inj_ ≤ 738 kPa) than stiffer biomaterials.

**Conclusions:** Combining the microstructural and biventricular finite element models enables quantifying micromechanics and signalling of transplanted cells in the heart. The approach offers a broader scope for *in silico* investigations of biomaterial and cell therapies for myocardial infarction and other cardiac pathologies.

## 1. Introduction

Cardiovascular disease-related deaths are predicted to rise from 17.5 million (i.e. 31% of global deaths) in 2012 to 22.2 million by 2030 [1, 2]. In low- and middle-income countries, such as South Africa, the number of affected people in the working-age population is increasing [3, 4]. Myocardial infarction (MI) causes cardiac cell death in the affected myocardial region, leading to scar formation, ventricular remodelling, and heart failure in the long term. Transplanting stem cells into the myocardium with injectable biomaterials has gained substantial interest in MI therapy research. The delivered stem cells are expected to promote regenerative reactions in the impaired tissue [5] and can differentiate into cardiac-like cells and endothelial cells (EC) necessary for revascularisation and functional recovery of the heart [6, 7].

Pre-clinical studies have demonstrated the beneficial effects of adult stem cell transplantation on MI. For example, Tomita et al. [8] demonstrated that intramyocardial transplantation of bone marrow-derived stem cells in the rat model improved angiogenesis and cardiac function and reported differentiation into cardiac-like cells (*in vitro* and *in vivo*) in the infarcted myocardium. Wang et al. [9] reported an increase in the expression of endothelial-related proteins, improved ejection fraction, reduced infarct size and a high vessel density by using a biochemically and genetically activated hydrogel loaded with adipose-derived stem cells in rats.

Clinical studies also reported the benefits of cell transplantation in MI. These benefits include the differentiation into cardiac-like cells [10-12] and enhanced neovascularisation and angiogenesis with reduced cardiomyocyte death through paracrine or autocrine signalling [13, 14]. Strauer and Steinhoff [15] observed improved cardiac function in patients with MI six months after human mesenchymal stem cell transplantation.

Pre-clinical and clinical studies support the hypothesis that paracrine signalling drives the beneficial effects of cell transplants in MI [8, 9, 13-19]. During cellular communication in cell-based therapies, proteins such as growth factors are released by a cell and attached to membrane receptors of the same or another cell through autocrine and paracrine effects. These processes are essential in the mechanotransduction of the injected cells in the dynamic mechanical environment of the heart.

Cyclic strain has been reported to play an essential role in the cellular production of growth factors and proteins that promote angiogenic phenotypes in bone marrow-derived mesenchymal stem cells [20, 21]. Zheng et al. [22] found an increase in vascular endothelial growth factor (VEGF) mRNA and its associated protein concentration in cardiac cells exposed to cyclic stretching. They also showed a 2.5-fold increase in transforming growth factor-beta (TGF-β) one-hour post-stretching. The beneficial effects of mechanically stimulated growth factor secretion in the infarcted heart were also demonstrated *in vivo* by several pre-clinical studies [23-25].

The physical environment of cells transplanted in the infarcted heart is essential in cell therapies. A biomaterial as a carrier and scaffold for the transplanted cells constitutes the immediate microenvironment that transmits mechanical stimuli from the myocardium to the cells. The biomaterial delivered into the infarct contributes to the mechanics of the surrounding myocardium and the embedded cells. Hence, the biomaterial offers a vehicle for improving wall mechanics and ventricular function and enhancing paracrine cardioprotective signalling through cellular mechanotransduction.

Several computational studies have investigated biomaterial delivery in the infarcted heart [26-31]. These studies focussed on determining the mechanical effects of the injected biomaterials on myocardial and ventricular mechanics that contribute to inhibiting post-infarction adverse remodelling. Wall et al. [29] reported a reduction in the wall stress and changes in ventricular function in the presence of an injected biomaterial that depended on the volume and stiffness of the delivered material and the delivery location using a finite element model of an ovine left ventricle. Other studies expanded on the investigation of treatment parameters to guide the optimisation of the injectate therapy using computational models of different species, including ovine [26, 30], canine [28], porcine [32, 33], human [34], and rat [31, 35, 36]. Wenk et al. [27] proposed an automated computational method to optimise injectate patterns for biomaterial delivery to treat heart failure, and Sirry et al. [35] was the first study presenting a microstructurally detailed *in situ* geometry of a hydrogel injectate in an infarcted rat heart. These studies exclusively focused on acellular biomaterial injectates, and there are no computational studies on the mechanics of the cells transplanted with the biomaterial into the infarcted heart.

Therefore, this computational study aimed to determine the mechanics of cells transplanted in a carrier biomaterial intramyocardially in a rat heart with a left ventricular infarct and quantify the relationship between the stiffness of the carrier biomaterial and the deformation of the embedded stem cells.

## 2. Materials and methods

The deformation of intramyocardially transplanted stem cells was investigated using a multi-scale approach linking the ventricular and wall mechanics to the cellular deformation (Figure 1). First, a biventricular finite element model of a rat heart with an infarct and intramyocardial biomaterial injectate in the left ventricle (LV) was developed and used to determine the ventricular and myocardial mechanics during a cardiac cycle (organ and tissue length scale). Second, a microstructurally detailed finite element model of a mid-wall region in the LV infarct with biomaterial injectate and transplanted therapeutic cells, referred to as the microstructural model, was developed (cellular length scale).

**Figure 1.**
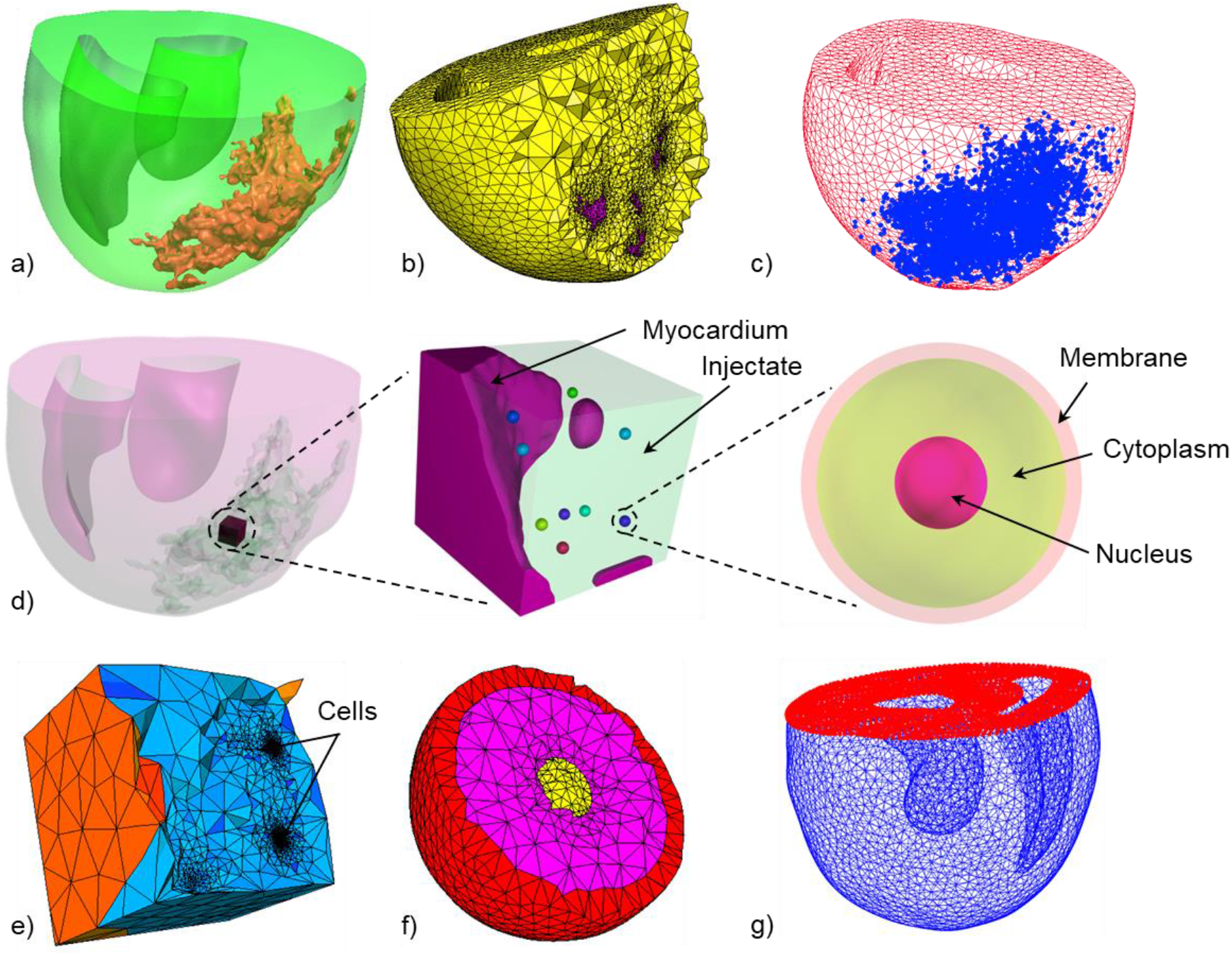
Geometries and models developed from the biventricular rat geometry to the microstructure extracted from the left ventricle mid-wall with transplanted cells in the injectate region. a) Biventricular geometry showing myocardium (green translucent) and injectate (beige). b) Meshed biventricular geometry with a cross-section illustrating the difference in mesh density between myocardium (yellow) and injectate (purple). c) Meshed biventricular geometry illustrating the infarct region nodes (blue). d) Development of a microstructural model of a left ventricle mid-wall region. Left: Biventricular geometry showing the size and location of the microstructural region. Middle: Microstructural geometry comprising myocardium, injectate and nine transplanted cells. Right: Cellular geometry with membrane, cytoplasm, and nucleus. e) Meshed microstructural geometry with coarse mesh for the myocardium and injectate and fine mesh for the cells. f) Cross-sectional view of meshed geometry of single cell showing membrane (red), cytoplasm (pink) and nucleus (yellow). g) The biventricular FE mesh’s basal nodes (red) were fixed as the boundary condition for simulations.

Nodal displacements of the LV mid-wall region of the microstructural geometry were recorded from the biventricular model during a cardiac cycle. The recorded displacements were applied as mechanical load boundary conditions to the outer surface of the microstructural model, resulting in deformation of the myocardium, injectate, and transplanted cells.

### 2.1 Geometrical modelling

#### 2.1.1 Biventricular geometry of an infarcted rat heart with biomaterial injectate

The current study utilised a three-dimensional biventricular geometry of a rat heart with infarct and polymeric intramyocardial injectate in the left ventricle (LV) developed in a previous study [36]. In brief, myocardial infarction was induced in a male Wistar rat by permanent ligation of the left anterior descending coronary artery and 100 µl of radiopaque silicone rubber containing lead chromate (Microfil® MV-120) was injected into the infarct region seven days later. The heart was explanted and underwent *ex vivo* microcomputed tomography (µCT) (custom scanner with Feinfocus X-ray tube and Varian 2520V Paxscan a-Si flat panel detector, Centre for X-ray Tomography, Ghent University, Belgium [38]). The biventricular geometry was reconstructed from the µCT image stack using semi-automated segmentation tools, including region-growing, level-set thresholding, and manual actions (Simpleware, Synopsys). The resulting geometry captured the essential morphology of the left and right ventricles and microstructural details of the dispersed injectate in the LV free wall (Figure 1 a). The meshed geometry comprised 147,240 (mesh density 302.8 mm^3^) and 58,902 (3,852.3 mm^3^) quadratic tetrahedral elements in the myocardial and injectate region, respectively (Figure 1 b). The meshed geometry was imported in Abaqus 6.14-3 CAE (Dassault Systèmes, Providence, RI, USA) and the infarct region was approximated by identifying the nodes surrounding the biomaterial injectate (Figure 1 c). The myofibre orientation varying from -50° at the epicardium to 80° at the endocardium [36] was implemented with a rule-based approach.

#### 2.1.2 Microstructural LV mid-wall geometry and transplanted cells

A microstructural geometry of mid-wall volume of 748 µm x 748 µm x 722 µm in the LV infarct (Figure 1 d) with higher spatial resolution was reconstructed by resampling of the cardiac µCT image data with reduced spacing (7.8 µm compared to 30 µm in x, y, and z direction for the biventricular geometry) (Simpleware).

Fifteen cells comprising membrane, cytoplasm and nucleus were numerically seeded at random locations in the injectate region of the microstructural geometry with a custom Python script in Simpleware ScanIP. For each cell, three concentric spherical surfaces with diameters of 60, 55, and 20 µm were created for the membrane, cytoplasm, and nucleus (Figure 1 d right). A Boolean subtraction resulted in a thickness of 5 µm and 35 µm for the membrane and the cytoplasm, respectively. The tree-component assembly was placed in the injectate. Of these 15 cells, six were located in the myocardium or near model boundaries and interfaces and were not considered for the analysis.

The resulting micro-structural geometry containing myocardium, injectate, and nine cells with membrane, cytoplasm and nucleus was meshed and imported into Abaqus. The mesh comprised 320,653 10-node tetrahedral elements (C3D10M), i.e. 2,552, 168,746, and 149,355 elements for the myocardium, injectate and the nine cells, respectively. The mesh size varied among the nine cells based on the automated meshing process (Simpleware), and element numbers were 6,101 ± 42 for the membrane, 8,710 ± 299 for the cytoplasm, and 1,784 ± 82 for the nucleus.

### 2.2 Constitutive laws

#### 2.2.1 Healthy and infarcted myocardium

The passive mechanical behaviour of the myocardium was represented by a hyperelastic anisotropic law using a modified strain energy density function from Holzapfel and Ogden [39], Eqn. (1). Changes were introduced to describe the pathological stage of the myocardium [40]. The passive mechanical response of infarcted myocardium depends on the stage of infarct healing [41-44]. A one-week infarct was modelled, corresponding to the delayed biomaterial delivery into the infarct. The associated increased stiffness in the fibre, circumferential, and longitudinal directions [45] was implemented using parameters h and p in Eqn. (2). The tissue health parameter 0 ≤ h ≤ 1 corresponds to infarcted (h = 0) and healthy (h = 1) myocardium [40]. The pathological scaling parameter p = 4.56 [40, 46] adjusts the passive material response of the myocardium according to pathology.

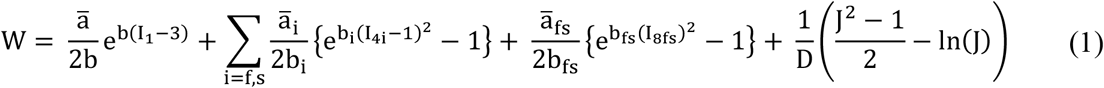

With

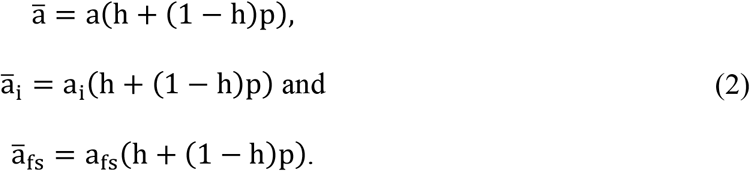

The active contraction of the myocardium was represented with a time-varying elastance approach with an additional term to consider the passive response during the contraction [47-49] and the tissue health parameter h described above [40].

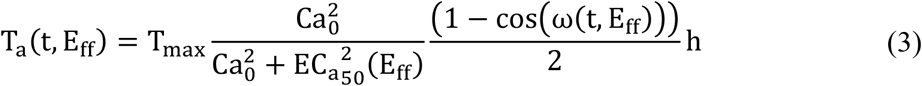

The time-varying elastance model was implemented by introducing an active tension-based force (T_a_), Eqn. <u>(3</u>), which depends on calcium concentration and the timing of the myocardial contraction, both driven by the sarcomere length [40, 49]. The active force function was multiplied by a health parameter, h (Eqn. <u>(3</u>)) to set the myocardium contractility to vary from 0 (fully infarcted myocardium) to 1 (healthy myocardium) [40].

The additive approach was used to compute the total tension in the myocardium where the time-varying active tension **T**_**a**_ in Eqn. <u>(3)</u>, was combined with the tension derived from the passive response **T**_**p**_:

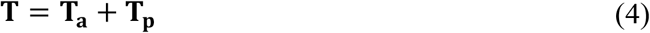

The active and passive constitutive parameter values were adopted from [40] and are provided in Supplemental Tables S1 and S2.

#### 2.2.2 Injectable biomaterial

The injectable biomaterial, e.g. a polyethylene glycol (PEG) hydrogel, was represented as a hyperelastic isotropic incompressible Neo-Hookean material:

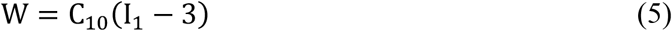

where I_1_ is the first deviatoric strain invariant, and C_10_ characterises the material stiffness obtained from the elastic modulus E_inj_ and the Poisson’s ratio ν_inj_:

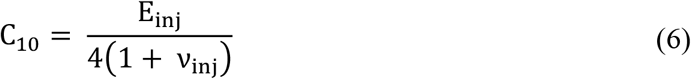

with ν_inj_ = 0.5 defining incompressibility and E_inj_ varied between 4.1 and 405,900 kPa (see section 2.4 FE simulations and data analysis for more details).

#### 2.2.3 Cells

Cell membrane, cytoplasm and nucleus were treated as hyperelastic isotropic compressible material using a Neo-Hookean strain energy density function [50, 51]:

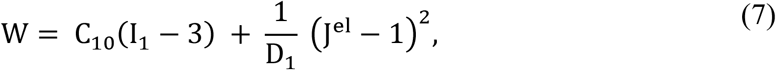

where J^el^ is a measure of volume change of the compressible material, and C_10_ and D_1_ depend on the elastic modulus and the material compressibility. The material parameter values for the cellular components are given in Supplemental Table S3.

### 2.3 Boundary conditions and loading

The boundary and loading conditions for the biventricular model to simulate a single cardiac cycle were described in detail previously [36]. In brief, zero displacement in the longitudinal direction was applied to the nodes at the basal surface to prevent the rigid body motion of the biventricular geometry (Figure 1 g). Additionally, a basal node in the LV ring was fixed in the circumferential and radial directions to prevent lateral motion and rotation. Passive filling was implemented with a linearly increasing pressure load on ventricular cavity surfaces from 0 mmHg to the end-diastolic pressure of 3.0 mmHg for the LV and 0.75 mmHg for the RV. Thereafter, the active contraction was initiated with the time-varying elastance representation of the myocardium (as described in section 2.2.1). The onset of the decline in active tension defined the end-systolic time point and the end of the cardiac cycle.

Displacement boundary conditions for the microstructural model were obtained from the biventricular model employing the node-based sub-modelling technique in Abaqus. The displacements of sets of nodes near the surfaces of the microstructural region were recorded in the biventricular geometry during the simulation of one cardiac cycle. These recorded displacements were applied as boundary conditions to surface nodes of the microstructural FE mesh and recapitulated the surface displacements and deformations during a cardiac cycle.

### 2.4 FE simulations and data analysis

A parametric study with the variation of the elastic modulus of the biomaterial injectate in the range of E_inj_ = 4.1 to 405,900 kPa (i.e. 4.1, 7.4, 40.6, 73.8, 405.9, 738, 4,059, 7,380, 40,590, 73,800 and 405,900 kPa) was conducted to determine the impact of biomaterial stiffness on the deformation of the transplanted cells in the biomaterial injectate in the infarct.

The analysis of the deformation of the transplanted cells comprised the following strain measures.

i. The element strain as mean strain at the integration points of an element:

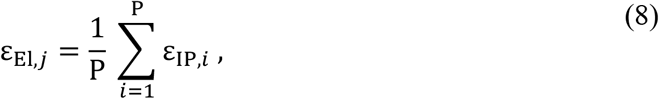

where ε_IP,*i*_ is the strain at integration point *i*, and P is the total number of integration points of element *j* in the cell mesh.
ii. The volume-averaged strain in a cell component (i.e. membrane, cytoplasm or nucleus) of cell *k*:

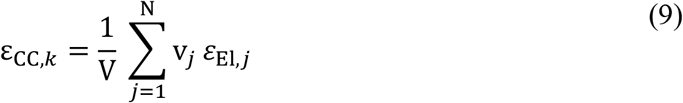

where V is the total volume of the cell component considered, N is the total number of elements in the cell component, v_*j*_ is the volume of element *j*, and ε_El,*j*_ is the element strain of element *j* calculated with Eqn. <u>(8)</u>.
iii. The mean strain in a cell component for all cells in the model:

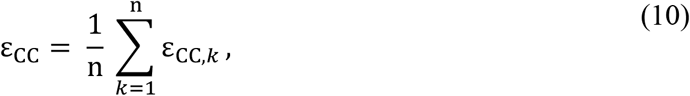

where ε_CC,*k*_ is the volume-averaged strain in a component of cell *k* according to Eqn. (9), and n is the total number of cells in the model (n = 9 in the current study).
iv. The mean strain in the entire cell for all n cells in the model also referred to as cell strain:

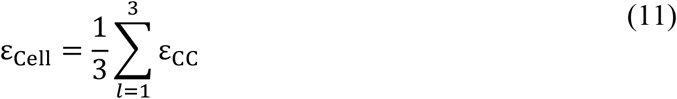

for the three cell components, i.e. membrane, cytoplasm, and nucleus, where ε_CC_ is the mean strain in a cell component for all cells in the model according to Eqn. (10).

Descriptive statistical analysis was performed on the strain data in the different model components to determine the normality (Shapiro-Wilk normality test) and variability (SciPy, https://scipy.org/ and NumPy, https://numpy.org/). Data are presented as mean and standard deviation if normally distributed; otherwise, they are presented as median and interquartile range.

## 3. Results

The end-systolic deformation and maximum principal strain of a cross-section of the microstructural model and the transplanted cells are illustrated in <u>Figure 2</u> for an injectate elastic modulus of E_inj_ = 7.4 kPa. The highest maximum principal strain was observed in the cellular membrane with the lowest elastic modulus (E = 1.7 kPa), whereas the strain in the cytoplasm (E = 8.0 kPa) and nucleus (E = 5.0 kPa) was similar to that in the biomaterial surrounding the cells.

**Figure 2.**
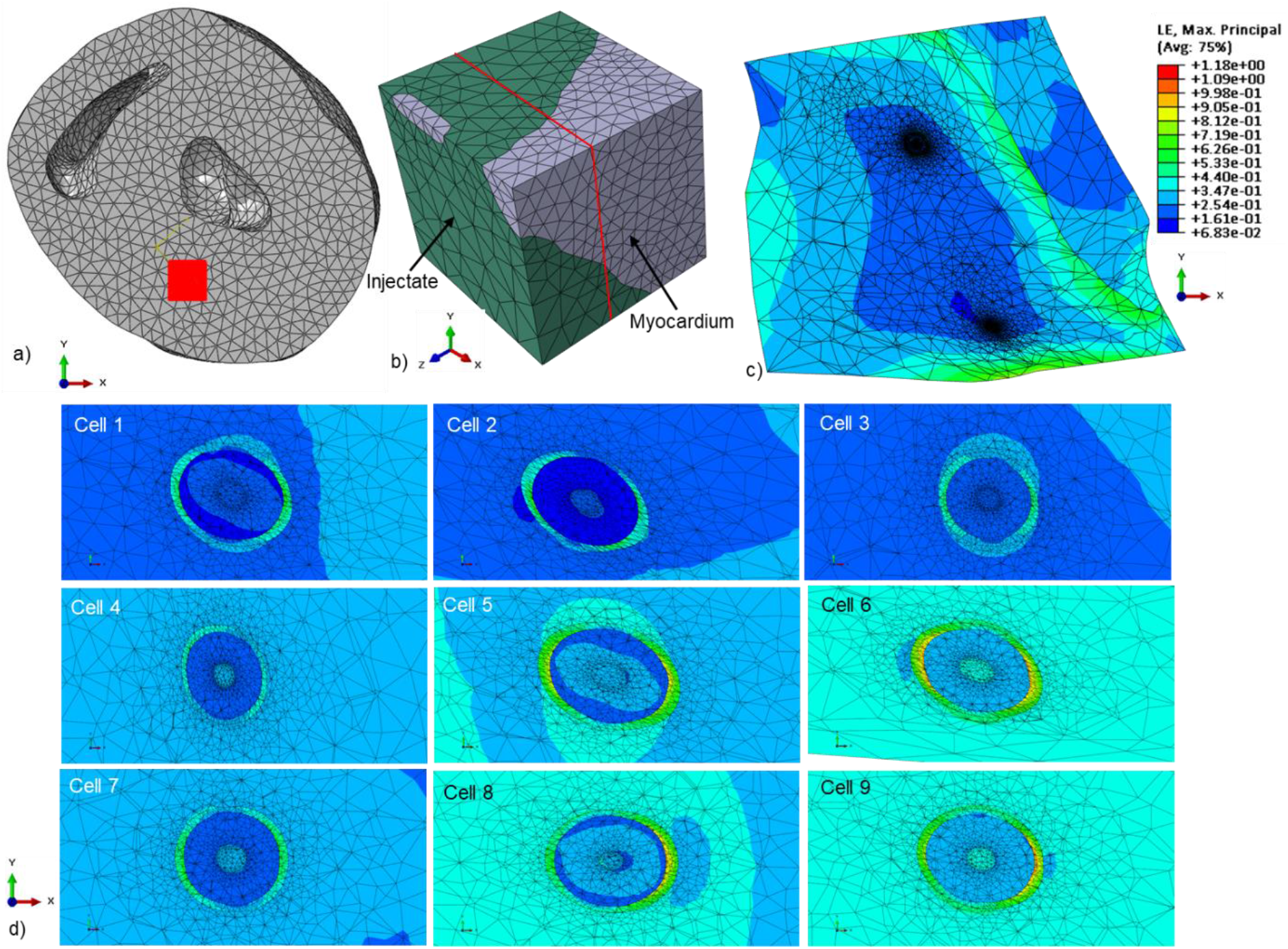
Illustration of deformation in infarcted myocardium, biomaterial injectate and transplanted cells at end-systole predicted with the microstructural FE model for E_inj_ = 7.4 kPa. a) Short-axis view of the undeformed biventricular geometry demonstrating position and orientation of the hexahedral region of the microstructural geometry in the LV free wall. b) Perspective view of the undeformed microstructural geometry containing infarcted myocardium and biomaterial injectate. c) Contour plot of maximum principal stress in a cross-section of the deformed end-systolic configuration of the microstructural model with colour legend of strain. The two regions with high mesh density are the locations of transplanted cells. (The location of the cross-section is indicated by red lines in b). d) Contour plots of maximum principal stress in a cross-section of the nine transplanted cells at different locations in the biomaterial injectate region of the microstructural model. The magnitude of the cellular deformation varied amongst the different cell locations. Generally, higher strain levels were predicted in the cellular membrane than in the cytoplasm and nucleus. The colour legend from figure c applies to these contour plots.

### 3.1 Deformation of injectate in the microstructural mid-wall model

At end-diastole, the median maximum principal strain in the injectate decreased from 6.5% to 0.9%, and the median minimum principal strain decreased in magnitude from -7.3% to -0.9% with increasing injectate elastic modulus from E_inj_ = 4.1 to 405,900 kPa (<u>Figure 3</u> a and b). At end-systole, the median maximum principal strain decreased from 43.8% to 1.4%, and the median minimum principal strain decreased in magnitude from -38.0% to -1.5% with increasing injectate elastic modulus (<u>Figure 3</u> c and d). The ratio of end-systolic to end-diastolic maximal principal strain decreased from 6.7 (for the softest injectate with E_inj_ = 4.1 kPa) to 1.6 (for the stiffest injectate with E_inj_ = 405,900 kPa). A similar decrease from 5.2 for the softest injectate to 1.6 for the stiffest injectate was observed for the ratio of end-systolic to end-diastolic minimum principal strain.

**Figure 3.**
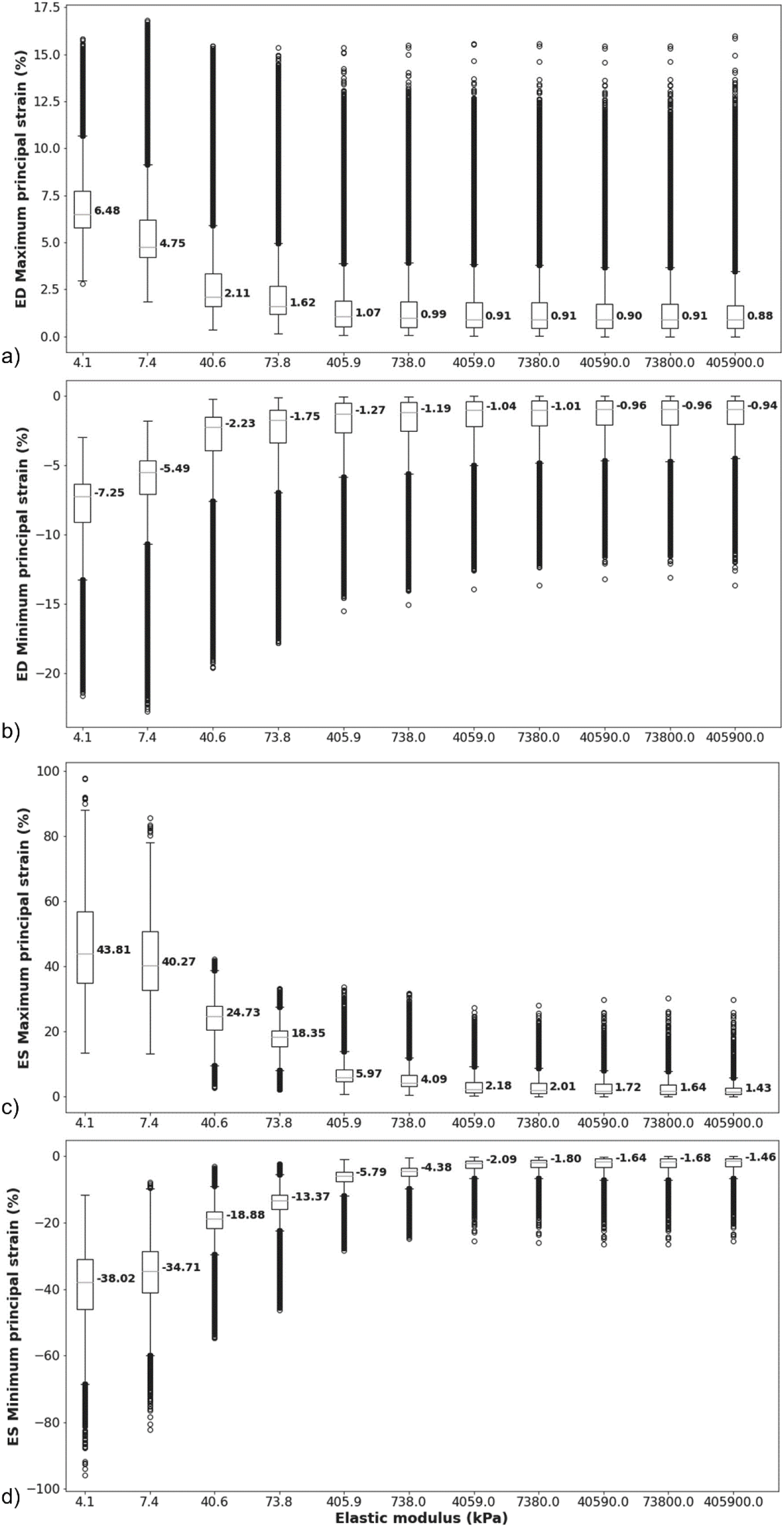
End-diastolic and end-systolic principal strains in the injectate of the microstructural model for different values of the injectate elastic modulus. Maximum (a) and minimum principal strain (b) at end-diastole, and maximum (c) and minimum principal strain (d) at end-systole. (Box and whiskers indicate the median (red line in box), interquartile range (IQR) between first and third quartile (lower and upper box bound), 1.5x IQR (lower and upper whisker), and data points smaller or larger than 1.5x IQR (open circles). Each data point represents the strain value ε_El_ in an element of the finite element mesh. Data larger or smaller than 1.5x IQR are considered actual data and not outliers.)

### 3.2 Deformation of transplanted cells in the injectate

The volume-averaged maximum principal strain in the cell components decreased for increasing injectate elastic modulus at end-systole and end-diastole (Figure 4 a-c). The strain decrease was larger in the lower range of the injectate elastic modulus from E_inj_ = 7.4 kPa to 738 kPa than for higher values of the injectate elastic modulus E_inj_ > 738 kPa. The membrane displayed higher strain than the cytoplasm and nucleus, which exhibited similar strains. Similar results were observed for the minimum principal strain, exhibiting negative values indicative of compression (<u>Figure 4 d-f</u>). The strain magnitude decreased for E_inj_ = 7.4 kPa to 738 kPa and levelled off for E_inj_ > 738 kPa. Increases in maximum and minimum principal strains at end-systole were observed for an increase of the injectate elastic modulus from E_inj_ = 4.1 kPa to 7.4 kPa.

**Figure 4.**
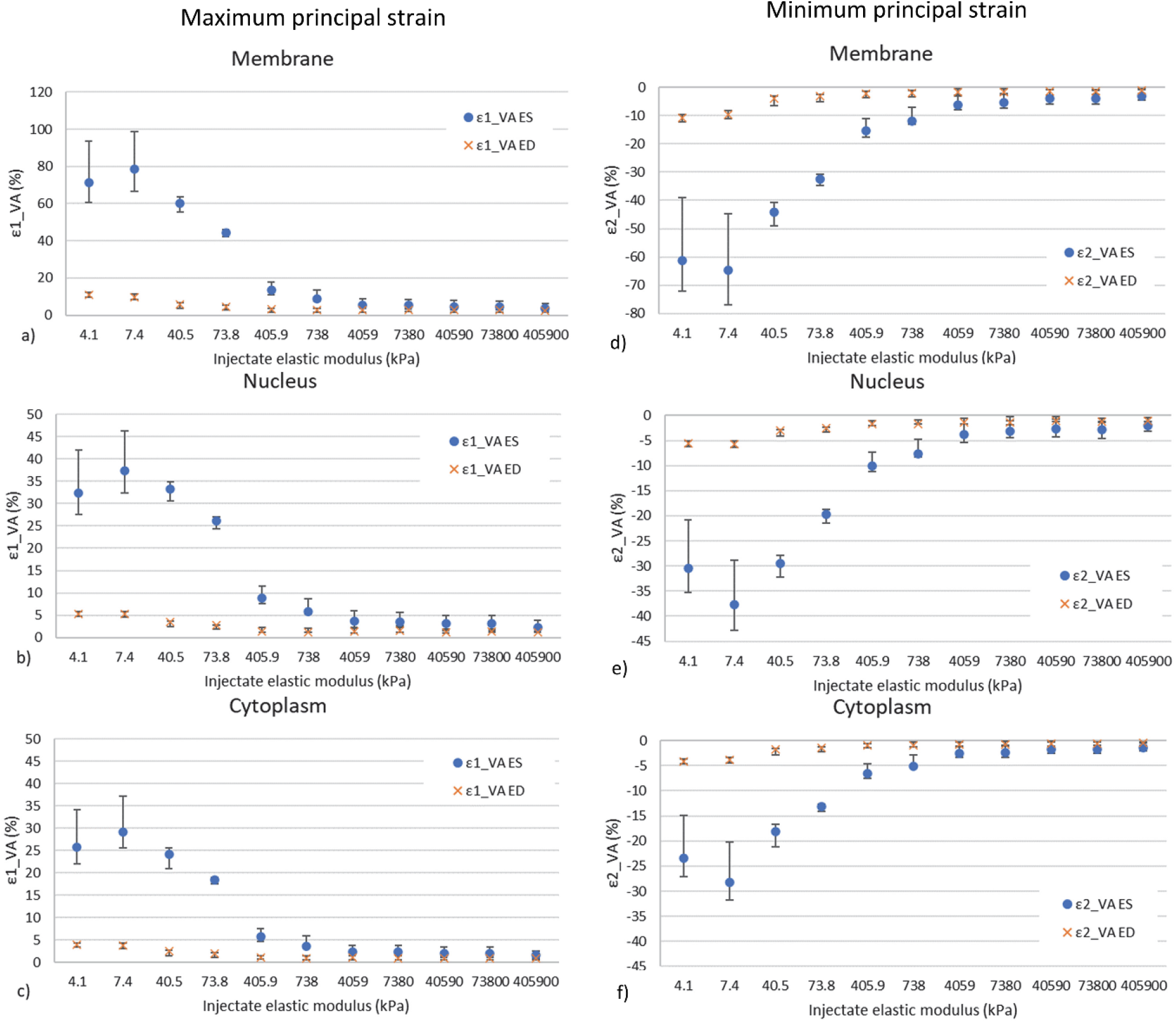
Maximum (a-c) and minimum (d-f) principal strains in the membrane, nucleus, and cytoplasm of transplanted cells at end-systole (ES) and end-diastole (ED) versus injectate elastic modulus. Data are median; error bars represent interquartile range.

The ES-ED range of the median volume-averaged maximum and minimum principal strains in the cell components decreased in magnitude for increasing injectate elastic modulus E_inj_ = 7.4 kPa to 405,900 kPa (Figure 5). The ES-ED range of the principal strain reflects the deformation range to which the transplanted cells are exposed during a cardiac cycle.

**Figure 5.**
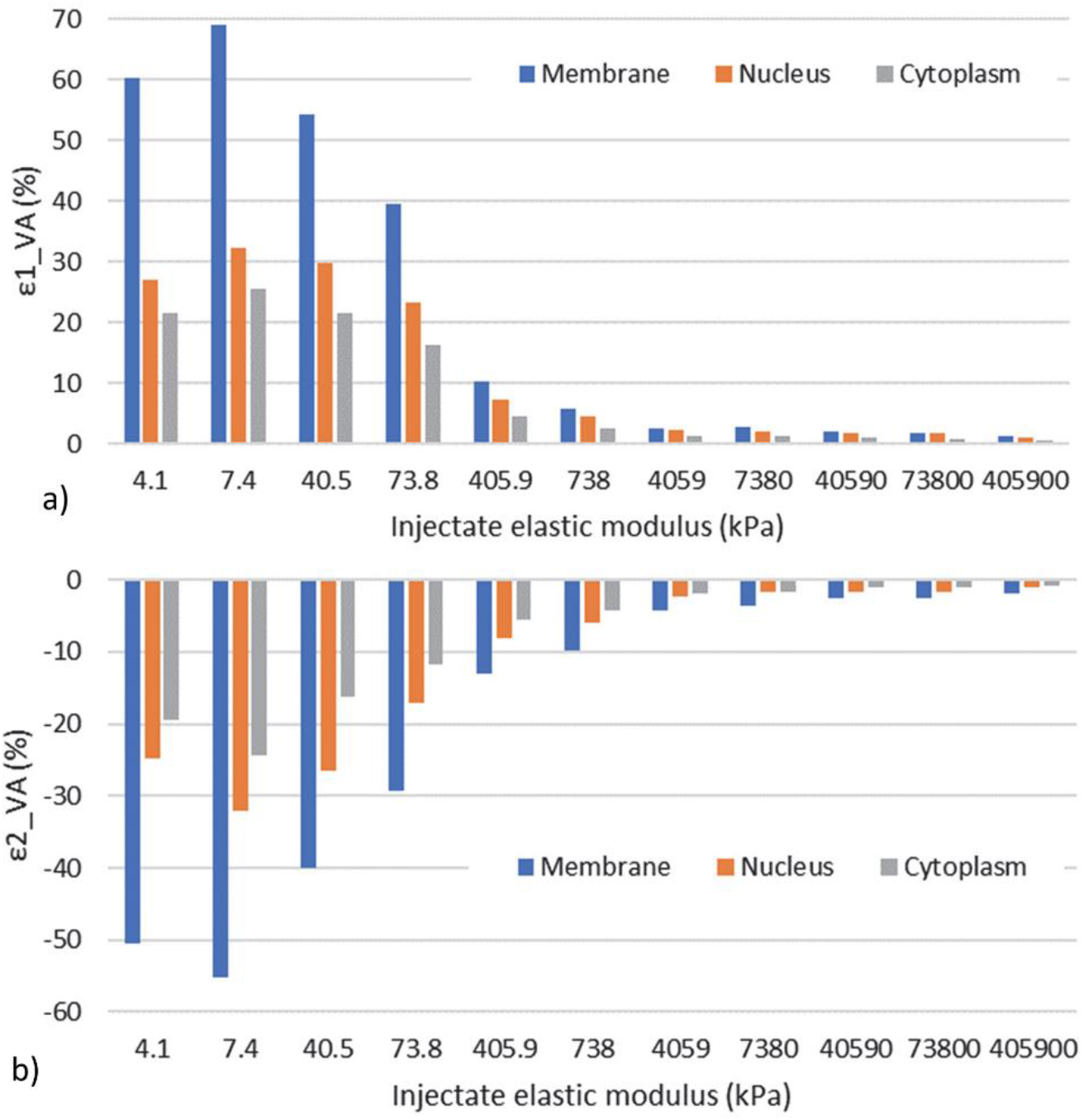
ES-ED range of maximum (a) and minimum (b) principal strain versus injectate elastic modulus for the three cell components.

## 4. Discussion

The present computational study demonstrated the effect of the stiffness of the biomaterial injectate on the mechanics of cells intramyocardially transplanted with the biomaterial. The variation of the injectate stiffness can be used to guide the deformation of the transplanted cells in the injectate. Soft injectates promise the most beneficial effects as the cell deformation is more sensitive to a change in injectate stiffness similar to the transplanted cells’ stiffness than higher injectate stiffnesses.

A decrease in strain in the injectate was predicted for an increasing injectate elastic modulus. Similar observations were reported by <u>Sirry et al. [35]</u>, where stiffer injectate resulted in less deformation within an acellular injectate region of the microstructural model. The current study investigated a wide range of injectate stiffness, with the upper range of the elastic modulus of E_inj_ > 738 kPa being unrealistic for PEG hydrogels and other injectable biomaterials [51, 52]. It was shown that the deformations in the injectate are negligible from a particular threshold of the injectate elastic modulus.

A decrease in ES and ED strains and the associated ES-ED strain range in the transplanted cells were predicted for increasing injectate stiffness. Additionally, the cell mechanics were considerably more sensitive to environmental stiffness changes for soft (E_inj_ ≤ 738 kPa) than stiff injectates (E_inj_ > 738 kPa).

Interestingly, an increase in cell strain was observed for an increase in injectate stiffness from E_inj_ = 4.1 kPa to 7.4 kPa. This contrasting behaviour may reveal a threshold of the injectate stiffness at which the mechanical response of the cells changes. This change may be explained when considering the elastic modulus of the cellular components, namely E = 1.7 kPa for the membrane, 5 kPa for the nucleus and 8 kPa for the cytoplasm. The injectate with E_inj_ = 4.1 kPa had a lower elastic modulus than the bulk of the cellular structure (i.e. cytoplasm and nucleus) and transferred less deformation to the cells than the injectate with E_inj_ = 7.4 kPa that more closely matches the elastic modulus of cytoplasm and nucleus.

The numerical cell seeding algorithm randomly placed cells in the microstructural model’s entire volume. Six of the 15 seeded cells were not used for the mechanical analysis due to their location in the tissue domain and near model boundaries and interfaces. Expanding the seeding algorithm to monitor the location of the cells will provide more control over the seeding process and the number of cells available for analysis. An increase in the number of seeded cells will allow a more detailed investigation of the impact of cell location and density on their mechanics. Such a comprehensive study can help design cell patterns within the injectate at the culture stage and when targeted spatial delivery into the heart becomes available.

The microstructural model used a single mesh configuration as the mesh variation was challenging due to the small thickness of the cellular membrane. Future studies may benefit from different meshing strategies or mesh sensitivity investigations. Using shell elements instead of three-dimensional elements for the cell membrane can be considered.

The cellular components were treated as isotropic and compressible materials with Neo-Hookean strain energy density functions. The constitutive equations did not consider the active processes in the cell (e.g. actin polymerisation and depolymerisation). Including these dynamic cytoskeletal processes will allow for assessing the medium- and long-term structural responses of the transplanted cells.

The biventricular FE model utilised structural boundary conditions at the truncated base as demonstrated in previous studies [32, 33, 36, 39]. Future studies can extend the biventricular model by including the atria and the pericardium in the geometry and representing the tissue environment of the heart in the chest in the boundary conditions. The current study was based on the simplifying assumption of the biventricular geometry reconstructed from μCT data representing the unloaded reference configuration for strain quantification. A validation of the model and predicted cardiac mechanics is needed for future studies. Furthermore, assessing the impact of the injectate stiffness on ED and ES time points may improve the model’s accuracy. A mesh sensitivity was not performed due to the detailed and realistic aspect of the reconstructed geometry.

The effect of the biomaterial injectate on the ventricular function of the infarcted heart has not been considered in the current study. However, functional improvement is essential for optimising biomaterial and stem cell-based therapy for MI and will be included in future work. While the developed biventricular model provides insights into the mechanical responses of cells to their environments, addressing the mentioned limitations may improve the accuracy of the predictions.

## 5. Conclusions

The current study is the first to quantify the deformation of therapeutic cells intramyocardially transplanted into an infarcted rat heart using a biomaterial injectate. The developed microstructural finite element model of the myocardium and biomaterial injectate at cellular length scale enables quantifying micromechanics of transplanted cells during a cardiac cycle. The coupled microstructural and biventricular cardiac finite element models can provide a point of departure for an *in silico* method including the mechanotransduction and signalling in the transplanted cells - with a broader scope of advancing therapeutic biomaterial and cell injections for MI and other cardiac conditions such as heart failure.

## Data availability

Computational models, custom code, and data supporting this study are available on the University of Cape Town’s institutional data repository (ZivaHub) at http://doi.org/10.25375/uct.22673884 as Motchon YD, Sack KL, Sirry MS, Nchejane NJ, Abdalrahman T, Nagawa J, Kruger M, Pauwels E, Van Loo D, De Muynck A, Van Hoorebeke L, Davies NH, Franz T. Data for “In silico mechanics and TGF-ß expression of stem cells intramyocardially transplanted with a biomaterial injectate for treatment of myocardial infarction”. ZivaHub, 2023, DOI: 10.25375/uct.22673884.

## Symbols

Symbol: Description
ā, b: Material parameter for the isotropic response in the myocardium
ā_fs_, b_fs_: Material parameter for the coupling stiffness in the fibre and sheet directions
ā_I_, b_i_: Material parameter for additional stiffness in the fibre and sheet directions for i = f, s
B: Governs the shape of peak isometric tension-sarcomere length relation
C_10_: Material parameter for the isotropic response in the Neo-Hookean model
Ca_0_: Peak intracellular calcium concentration
D: Myocardium material parameter for incompressibility properties
D_1_: Material parameter for incompressibility properties in the Neo-Hookean models
E: Elastic modulus for the isotropic responses in the Neo-Hookean model
ECa_50_: Length-dependent calcium sensitivity
F: Deformation gradient tensor
h: Parameter defining the pathological state of the tissue (healthy or infarcted)
I_4f_: Transversely isotropic invariant in the fibre direction
I_4s_: Transversely isotropic invariant in the sheet direction
I_8fs_: Orthotropic invariant from coupling in fibre and sheet direction
I_i_: Isotropic invariants in principal directions
J: Third deformation gradient invariant describing the volume change of a compressible material
l: Sarcomere length (time-varying model in active contraction)
l_r_: Initial sarcomere length (time-varying model in active contraction)
p: Parameter scaling the isotropic response of the infarcted tissue
T: Stress tensor defining the mechanics of the myocardium
T_a_: Active stress tensor describing the myocardium responses during active contraction
T_max_: Constitutive law scaling factor (time-varying model in active contraction)
T_p_: Passive stress tensor defining the additional stress from the passive response during the active contraction
t: Time
t_r_: Linear function of sarcomere length
t_0_: Time to reach peak tension (time-varying model in active contraction)
W: Strain energy density function
ε: _Cell_Cell strain
ε_Sub_: Substrate strain
κ: Bulk modulus
λ: Stretch
ν: Poisson’s ratio
ω: Variable in the internal variable function depending on time and sarcomere length
σ: Stress

## CRediT Author Contributions

YDM: Data curation, Formal analysis, Investigation, Methodology, Project administration, Software, Validation, Visualization, Writing – Original Draft Preparation, Writing – Review & Editing

KLS: Methodology, Software, Supervision, Writing – Review & Editing

MSS: Methodology, Software, Supervision, Writing – Review & Editing

NJN: Methodology, Writing – Review & Editing

TA: Methodology, Supervision, Writing – Review & Editing

JN: Methodology, Writing – Review & Editing

MK: Investigation, Writing – Review & Editing

EP: Investigation, Methodology, Writing – Review & Editing

DVL: Investigation, Methodology, Writing – Review & Editing

ADM: Investigation, Writing – Review & Editing

LVH: Resources, Methodology, Supervision, Writing – Review & Editing

NHD: Conceptualization, Data curation, Funding acquisition, Methodology, Project administration, Resources, Supervision, Validation, Writing – Review & Editing

TF: Conceptualization, Data curation, Funding acquisition, Methodology, Project administration, Resources, Supervision, Validation, Writing – Original Draft Preparation, Writing – Review & Editing

