## Supplemental tables for "*In silico* mechanics of stem cells intramyocardially transplanted with a biomaterial injectate for treatment of myocardial infarction"

**Supplemental Material**

Table S1. Constitutive parameters for passive mechanical behaviour of myocardium (Sack et al. 2018)

| Parameter | Value & unit | Description |
| --- | --- | --- |
| $\bar{\mathbf{a}}$ | a = 0.2065 kPa | Governs the isotropic response |
| ${\bar{\mathbf{a}}}_{\mathbf{i}}$ | a_f_ = 0.68 kPa  a_s_ = 0.945 kPa | Governs stiffness in the fibre (a_f_) and sheet (a_s_) direction for healthy myocardium, for h = 1 |
|  | Directly proportional to a_f_, a_s_ | Governs additional stiffness in the fibre (a_f_) and sheet (a_s_) direction for infarcted myocardium, for h = 0. It is scaled by the parameter p |
| ${\bar{\mathbf{a}}}_{\mathbf{fs}}$ | a_fs_ = 5.55 x 10^-2^ kPa | Governs coupling stiffness in the fibre and sheet direction for healthy myocardium |
|  | - | Governs coupling stiffness in the fibre and sheet direction for infarcted myocardium |
| b | 7.61 | Governs the isotropic response |
| b_f_ | 14.61 | Governs additional stiffness in the fibre direction |
| b_s_ | 12.67 | Governs additional stiffness in the sheet direction |
| b_fs_ | 3.12 | Governs coupling stiffness in the fibre and sheet direction |
| I_1_, I_4i_, I_8fs_ | - | Strain invariants |
| D | 0.2 x 10^-3^ kPa^-1^ | Defines the material incompressibility. Inversely proportional to the bulk modulus (κ i.e. κ=2/D), defining the material’s resistance to compression) |
| h | [0, 1] | Governs the health of the myocardium material point (h = 0 or 1 for infarcted or healthy myocardium) |
| p | 4.56 | Adjusts the passive response according to the stage of the infarct |
| J | - | Third deformation gradient invariant |

Table S2. Constitutive parameters for active contraction in the myocardium (Guccione et al. 1993; Sack et al. 2018)

| Parameter | Value & unit | Description |
| --- | --- | --- |
| T_max_ | 60 kPa | Constitutive law scaling factor |
| Ca_0_ | 4.35 µmol/l | Peak intercellular calcium concentration |
| Ca_0max_ | 4.35 µmol/l | Maximum intercellular calcium concentration |
| B | 4,750 mm^-1^ | Governs the shape of the peak isometric tension-sarcomere length relation |
| l_0_ | 1.58 x 10^-3^ mm | Sarcomere length below which no active force develops |
| t_0_ | 0.15 s | Time to reach peak tension |
| m | 1,048.9 s/mm | Govern the shape of the linear relaxation duration and sarcomere relaxation |
| b | 1.5 s | Govern the shape of the linear relaxation duration and sarcomere relaxation |
| E_ff_ | - | Lagrangian strain tensor component aligned with the local muscle fibre direction |
| l_r_ | 2.03 x 10^-3^ mm | Initial sarcomere length |

Table S3. Values of mechanical properties of cell components (Caille et al. 2002; Jean et al. 2005)

|  | Elastic modulus E (kPa) | Poisson ratio ν (-) |
| --- | --- | --- |
| Membrane | 1.7 | 0.4 |
| Cytoplasm | 8 | 0.3 |
| Nucleus | 5 | 0.4 |

Table S4. Material parameters and values for components of cell stretching FE model (Rodriguez et al. 2013)

|  | **Substrate** | **Focal adhesion** | **Cell** |
| --- | --- | --- | --- |
| Volumetric mass density ρ (kg/m^3^) | - | - | 1,060 |
| Elastic modulus E (MPa) | 1 | 0.05 | - |
| Poisson ratio ν (-) | 0.49 | 0.3 | 0.3 |
| C_10_ (MPa) | - | - | 0.00154 |
| D_1_ (MPa) | - | - | 300 |

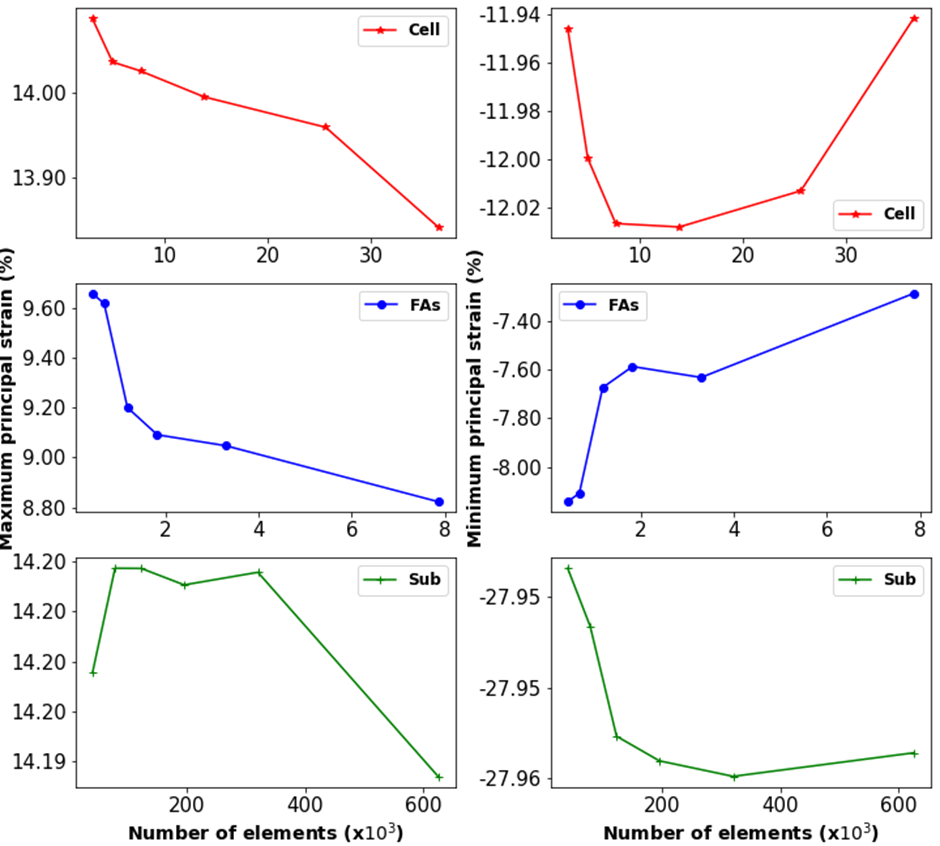

Figure S1. Mesh sensitivity study for the single-cell stretching finite element model. Convergence of the maximum (left column) and minimum (right column) principal strain for increasing number of elements in the cell body (top), focal adhesions (middle) and substrate geometries (bottom) for a biaxial substrate strain of 15%. Based on this study final mesh sizes were 7,697 for the cell body, 1,186 for each FA disk, and 122,694 for the substrate.
